# Bacteriophage deficiency characterizes respiratory virome dysbiosis in childhood asthma

**DOI:** 10.1101/2020.08.04.236067

**Authors:** Spyridon Megremis, Bede Constantinides, Paraskevi Xepapadaki, Claus Bachert, Susetta Neurath-Finotto, Tuomas Jartti, Marek L Kowalski, Alexandros Georgios Sotiropoulos, Avraam Tapinos, Tytti Vuorinen, Evangelos Andreakos, David Robertson, Nikolaos Papadopoulos

## Abstract

Asthma development and persistence is tightly linked to respiratory viruses. Viral presence is usually interrogated with targeted approaches during periods of disease activity and/or infections, thus neglecting viral occurrence during steady states. We investigate the virome in the upper respiratory system of healthy and asthmatic preschool children during asymptomatic/non-infection periods using metagenomics. Children with asthma have a characteristically dysbiotic virome that correlates to disease severity and control. The major component of dysbiosis is bacteriophage deficiency, while eukaryotic viral presence is increased. At the metacommunity level, differential virus species co-occurrence patterns suggest a decrease of the microbiota community resilience in asthma. Viral dysbiosis is therefore a key characteristic of asthma pathophysiology.

## Introduction

More than 300 million people suffer from asthma worldwide^1^, affecting their quality of life, and reducing educational attainment in paediatric and adolescent populations^2^. Asthma remains a major driver of morbidity throughout the productive years, while also affecting healthy aging^3^. Asthma susceptibility to viral and bacterial respiratory infections may influence the divergence from a healthy trajectory and is one of the main priorities for research and intervention ^1,4-8^.

The respiratory tract harbours heterogenous microbiota decreasing in biomass and richness from the ‘source’ upper respiratory tract (URT) towards the lower airway based on the microbial immigration and elimination model^9,10^. It has been demonstrated that bacterial communities are disrupted in disease or infectious conditions^11-15^, while the bacterial component of the respiratory microbiome is increasingly recognised to have an important role in the susceptibility and severity of acute respiratory illness and asthma^16,17^. The microbial colonisation of the nasal cavity and nasopharynx in early life has been linked with wheeze episodes and progression to asthma, potentially mediated through resistance or susceptibility to acute respiratory infections during childhood^18-22^.

Compared to cellular microbes, respiratory viruses are currently considered the most important drivers of asthma development, exacerbation and persistence^23-27^. Upper respiratory viruses, most importantly rhinoviruses (RV), are strongly implicated in the induction of airway hyperresponsiveness and airway remodelling^1,4,28-31^. Impairment of innate immune responses in children with asthma results in defective pathogen recognition, impaired interferon release and suboptimal antiviral responses^32,33^, and drives the development of biased T2 inflammation^32,33^.

Notably, acute respiratory viruses have historically been evaluated in isolation from the respiratory microbial ecosystem^34^. Most importantly, our knowledge on prokaryotic viruses infecting bacteria (bacteriophages or phages) is extremely limited, even though they are the most straightforward link between viruses and bacteria in the respiratory system^35^. Only a handful of studies have investigated the respiratory prokaryotic virome^36,37^ and to our knowledge none in asthma.

We hypothesized that a respiratory virome, including both eukaryotic and prokaryotic viruses, is present in the upper airway and is affected in asthma. The aim of this study was to identify DNA and RNA virus species in the upper respiratory system of healthy and asthmatic preschool children during a period of stable disease activity using a metagenomic approach^38^. We provide key characteristics of viral dysbiosis in asthma, in the context of the respiratory virome.

## Results

### Reduced bacteriophage presence in asthma

We processed nasopharyngeal samples (NPS) obtained from a subgroup of children with asthma and healthy controls recruited in the PreDicta cohort ^27^ (Figure S1 & S2A). The children did not have symptoms of a respiratory infection or asthma exacerbation for at least a month prior to sampling. Subject characteristics are shown in Supplementary table 1. The viral metagenome assembled genomes (vMAGs) identified in the discovery cohort (Figure S2B-S2D) were taxonomically organised in the order of the prokaryotic Caudovirales viruses (phages) and eukaryotic viruses of the Anelloviridae and Picornaviridae families. The phage genome frequency of occurrence was significantly less in asthma (p:0.045) (Figure 1A). We observed that 87.5% (7 out of 8) of these vMAGs were identified in healthy samples compared to 37.5% (3 out of 8) in asthma (Figure 1A), along with an elevated number of sequencing reads mapping to them (Figure 1B). We estimated the number of prokaryotic vMAGs as a faction of increasing sample size and observed a clear divergence between the two groups (Figure 1C). Further sample-level metagenomic analysis revealed that phages were significantly underrepresented in asthma (p:0.02) (Figure 1D).

**Figure 1:**
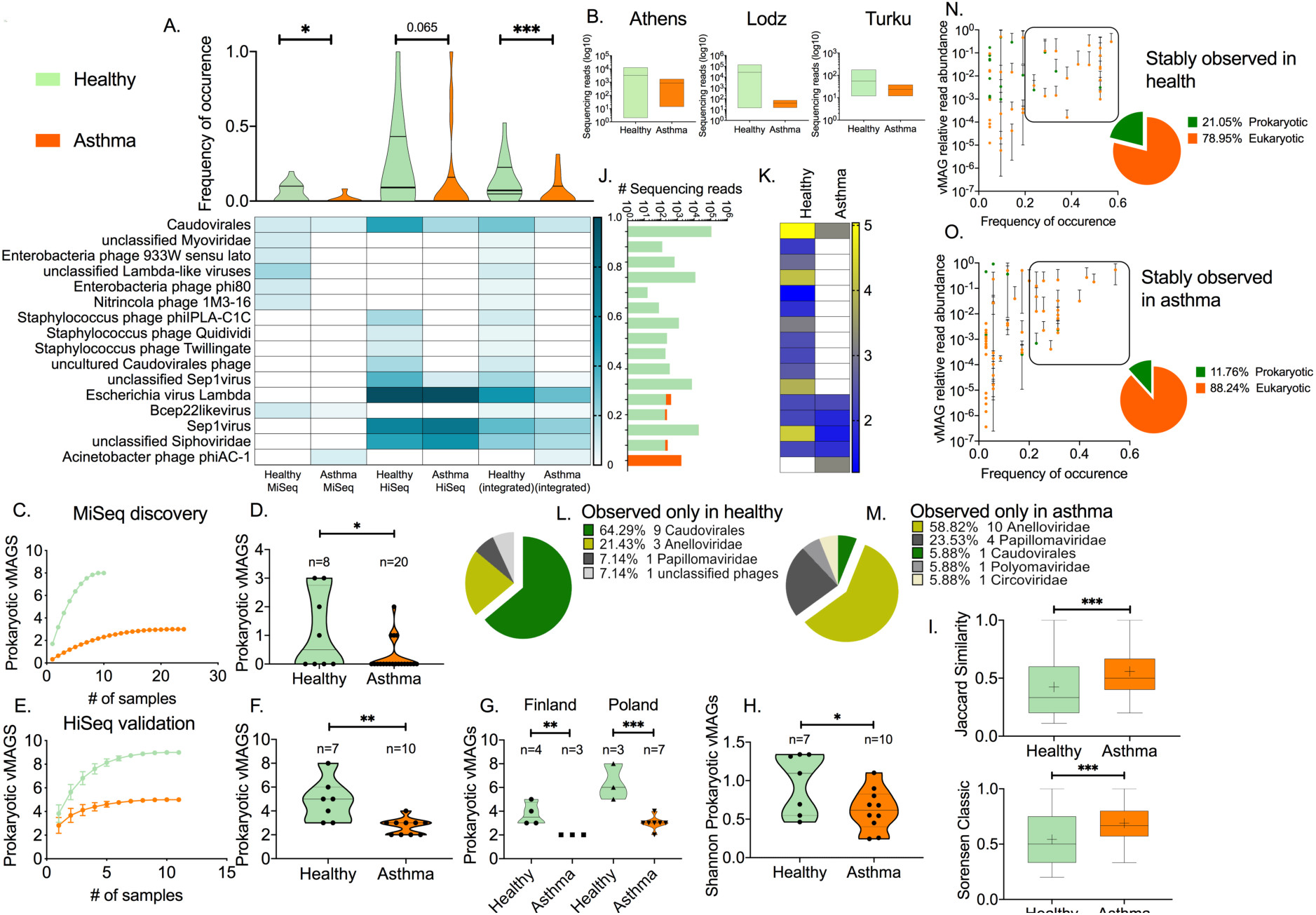
Prokaryotic viruses in health and asthma. (A) Heatmap of the frequency of occurrence (FrO) for each prokaryotic vMAG; Data on the discovery MiSeq and validation HiSeq cohorts are presented, along with the total vMAG FrO per clinical group (integrated data). FrO distributions are depicted as violin plots along with the median and 95%CIs. Statistical differences between viral species in health and asthma were tested using paired two-tailed t test. (B) Floating bar plots of the total number of sequencing reads that mapped to each prokaryotic vMAGs in health and asthma in Greece, Poland and Finland; Mean values and range are depicted. (C) Comparison of sample-based estimated number of prokaryotic vMAGs in health and asthma; MiSeq discovery cohort. The graph depicts the accumulation of bacteriophage vMAGs with increasing number of samples after randomisation without replacement. Data points depict mean values calculated after 100 repetitions with standard deviations. (D) Scatter plots of the number of prokaryotic vMAGs observed in each sample in health and asthma. Median and 95%CIs are depicted. Statistical significance was tested using a parametric two-tailed t test. (E) Comparison of sample-based estimated number of prokaryotic vMAGs in health and asthma; HiSeq validation cohort. Data points depict mean values calculated after 100 repetitions with standard deviations. (F) Comparison of the prokaryotic vMAG richness between healthy and asthmatic individuals; HiSeq cohort. Median and 95%CIs are depicted. Statistical significance was tested using a parametric two-tailed t test. (G) Comparison of the prokaryotic vMAG richness between healthy and asthmatic individuals in Poland and Finland; HiSeq cohort. Median and 95%CIs are depicted. Statistical significance was tested using a parametric two-tailed t test. (H) Comparison of the prokaryotic vMAG Shannon diversity between healthy and asthmatic individuals; HiSeq cohort. Median and 95%CIs are depicted. Statistical significance was tested using a parametric two-tailed t test. (I) Prokaryotic vMAG beta diversity within health and asthma. Floating bars depict range, mean (cross) and median of the Jaccard and Sorensen indexes calculate between the samples within each group; HiSeq cohort. Statistical significance was tested using a nonparametric Mann Whitney test. (J) Stacked plot of the fraction of sequencing reads aligned onto prokaryotic vMAGs identified in health and/or asthma (Log10 scale). Prokaryotic vMAGs are presented based on Figure 1A. (K) Heatmap of the total number of prokaryotic vMAG sequencing reads in health and asthma; scale (Log10); gradient: yellow-high to blue-low. Prokaryotic vMAGs are presented based on Figure 1A. (L) Compositional pie charts of viral genomes observed exclusively in healthy children. (M) Compositional pie charts of viral genomes observed exclusively in children with asthma. Scatter plots of vMAG frequency of occurrence and vMAG mean relative read abundance in (N) health and (O) asthma: We defined stably observed viruses based on their average sequencing read abundance and frequency of occurrence; the proportion of prokaryotic vMAGs within this subset was higher in health compared to asthma. Colour-coding: Orange; asthma Green; health. Significance tests: * p<0.05, ** p<0.001, *** p<0.0001, **** p<0.00001.

To validate our findings, we repeated the comparison in a second cohort of age-matched healthy and asthmatic children with increased Illumina read length and depth (validation cohort, Supplementary table 1) (Figure S2A). Compared to the discovery MiSeq output, this approach increased the number of contigs 5-fold, the number of vMAGs 2-fold, the viral sequencing reads 22-fold, while including samples from different geographical locations (Figure S2A, S2B & S2E). As in the discovery cohort, the identified prokaryotic viruses belonged exclusively to the Caudovirales order of phages (Figure S2F). Most eukaryotic viral contigs mapped to acute respiratory viruses of the Picornaviridae (Rhinoviruses), Paramyxoviridae (Parainfluenza) and Anelloviridae vMAGs (Figure S2F). There was no significant difference in the prokaryotic vMAG frequency of occurrence (p:0.065) (Figure 1A). However, similar to the discovery cohort, a lower number phage vMAGs were observed in asthma (55.5%: 5/9) compared to health (100%: 9/9) (Figure 1A), along with an elevated number of sequencing reads per vMAG, regardless of geography (Figure 1B). The health and asthma sample-based rarefaction curves diverged showing reduced number of different phages in asthma (Figure 1E). The reduction was also clearly present independent of the geographical location, confirming a generalised observation (Figure S3). To describe phage ecology within each sample, we focused on samples with two or more prokaryotic vMAGs: The prokaryotic viral richness was significantly reduced in asthma compared to healthy controls (p:0.003) (Figure 1F). This was also observed when the geographical origin of the samples was considered (Figure 1G). The within-sample Shannon diversity of prokaryotic viruses was significantly decreased in asthma (p:0.045) (Figure 1H). Asthmatics were more similar to each other compared to healthy controls based on the prokaryotic virome unweighted composition (presence/absence), measured using the Jaccard (p:0.0008) and Sorensen (p:0.0008) indices of beta diversity (Figure 1I).

Overall, the majority of the prokaryotic sequencing reads were observed in samples from healthy children (Figure 1J). Furthermore, there were fewer sequencing reads in asthma for most of the identified phage vMAGs (n=12, 75%) (Figure 1K). Phages were also over-represented in the subset of vMAGs identified only in healthy children (n=9, 64.29%) compared to the asthma-specific viral genomes (n=1, 5.88%) (Figure 1L & 1M). Within the ‘core’ virome, defined as vMAGs with more than 30% detection rate and more than 0.01% read abundance, bacteriophages accounted for 21% in healthy samples (Figure 1N), in contrast to 12% in asthma (Figure 1O). Collectively, these data show that the virome of preschool children with asthma is less rich and diverse in bacteriophages compared to healthy controls. This was observed in samples from three different geographical locations and at two different sequencing depths.

### Increased eukaryotic virus presence in asthma

We then asked whether eukaryotic viruses followed a similar trend. 14/15 (93%) eukaryotic vMAGs were observed in the asthma samples, versus 60% (n=9) in samples from the healthy group (Figure 2A & 2B). The frequency of occurrence of eukaryotic vMAGs was higher in asthma in the HiSeq cohort (p:0.016) and the integrated dataset (p:0.002). The healthy and asthmatic sample-based rarefaction curves of the estimated number of eukaryotic vMAGs diverged in both the discovery and validation cohorts demonstrating increased eukaryotic vMAG presence in asthma (Figure 2C & 2D), regardless of geography (Figure S3). There were no significant differences when considering eukaryotic vMAG presence within each sample, nevertheless, in the discovery cohort we observed a marginal increase (p: 0.059) in the abundance-based Shannon diversity (Figure 2E) and a significant (p: 0.026) reduction in the Simpson evenness of asthma samples (Figure 2G). These differences were also present in the validation cohort when comparing healthy and asthma samples from Poland: Shannon diversity was significantly increased (p: 0.045) (Figure 2F) while the Simpson evenness was significantly decreased in asthma (p: 0.029) (Figure 2H). This however was not observed in samples from Finland. Collectively, these data suggest that the asthmatic virome is often more diverse in eukaryotic viruses which acquire a more even distribution within each individual’s virome compared to the healthy viromes.

**Figure 2:**
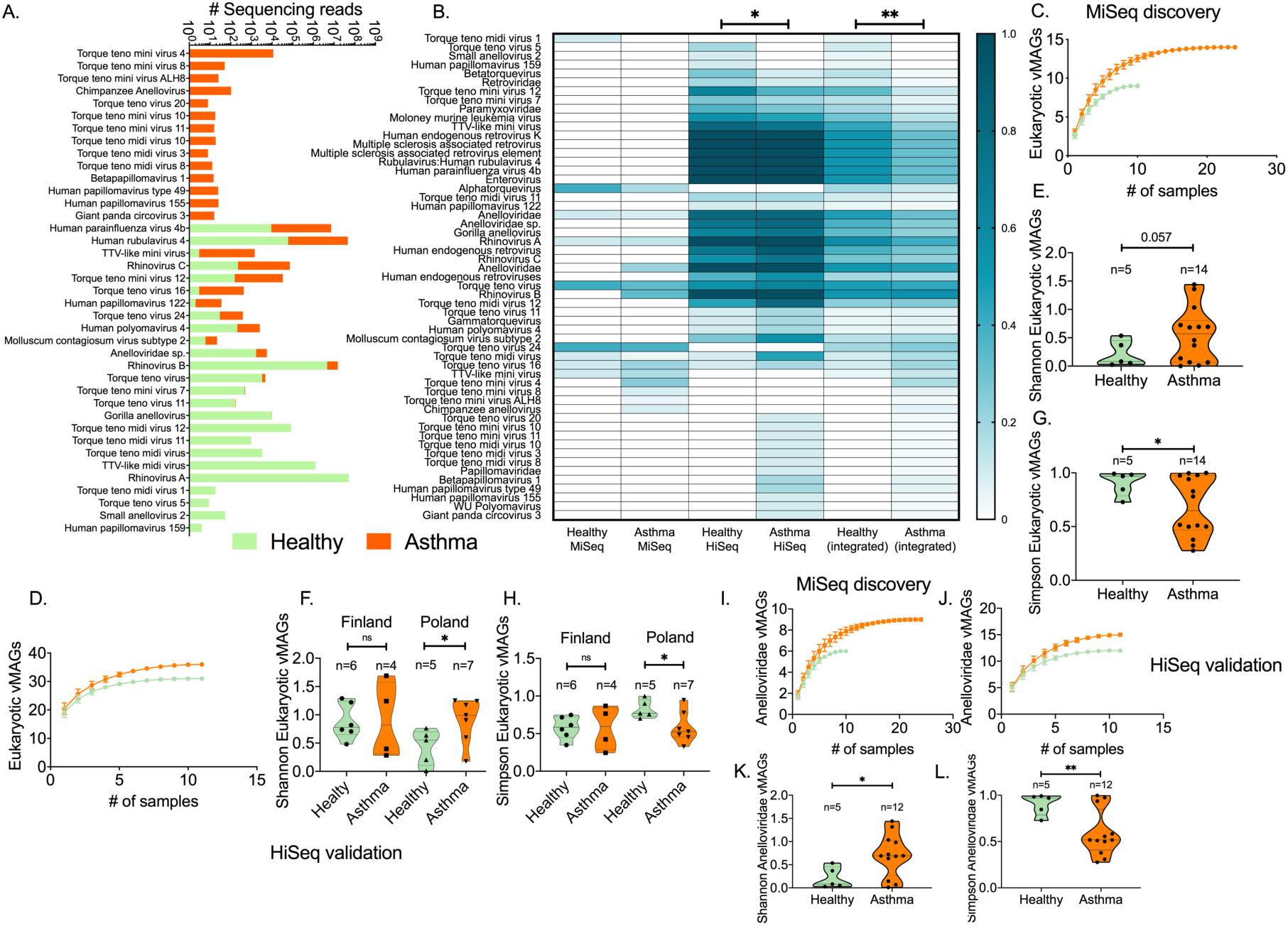
Eukaryotic viruses in the healthy and asthmatic virome. (A) Stacked plot of the fraction of sequencing reads aligned onto eukaryotic vMAGs identified in health and/or asthma (Log10 scale). (B) Heatmap of the frequency of occurrence (FrO) for each eukaryotic vMAG; Data on the MiSeq and HiSeq cohorts are presented, along with the FrO per clinical group (integrated dataset). Significant differences were tested using a two-tailed paired t test. Comparison of sample-based rarefaction curves of the estimated number of eukaryotic vMAGs in health and asthma; (C) MiSeq discovery, and (D) HiSeq validation cohort. Data points depict mean values calculated after 100 repetitions with standard deviations. Comparison of the eukaryotic vMAG Shannon diversity between healthy and asthmatic individuals; (E) MiSeq discovery, and (F) HiSeq validation cohort. Median and 95%CIs are depicted. Statistical significance was tested using a parametric two-tailed t test with Welch’s correction. Comparison of the eukaryotic vMAG Simpson evenness between healthy and asthmatic individuals; (G) MiSeq discovery, and (H) HiSeq validation cohort. Median and 95%CIs are depicted. Statistical significance was tested using a parametric two-tailed t test with Welch’s correction. Comparison of sample-based rarefaction curves of the estimated number of Anelloviridae vMAGs in health and asthma; (I) MiSeq discovery, and (J) HiSeq validation cohort. Data points depict mean values calculated after 100 repetitions with standard deviations. Comparison of the Anelloviridae vMAG ecological indexes between healthy and asthmatic individuals; (K) Shannon diversity, and (L) Simpson evenness. Median and 95%CIs are depicted. Statistical significance was tested using a parametric two-tailed t test with Welch’s correction. Colour-coding: Orange; asthma Green; health. Significance tests: * p<0.05, ** p<0.001, *** p<0.0001, **** p<0.00001.

### Increased presence of Anelloviruses in asthma

Anelloviruses accounted for the majority of eukaryotic vMAGs in both health (88.8%) and asthma (85.7%) in the MiSeq cohort and more than 45% in the HiSeq cohort (48.5% and 46.5%, respectively) (Figure 2A & 2B), representing a significant fraction of the respiratory eukaryotic virome (Figure S2D & S2F). Anelloviruses were the most frequent eukaryotic virus identified in the discovery cohort as well as in the validation cohort (Figure 2B). Most of the identified Anelloviridae genomes were observed in asthma (Figure 2A & 2B). Anelloviruses were enriched (14/20 of Anelloviridae vMAGs) among the eukaryotic vMAGs which were observed with higher frequency in asthmatics compared to healthy donors (Figure 2B), as well as among the vMAGs observed only in asthma (Figure 1M). The estimated number of Anelloviridae vMAGs as a fraction of sample size was higher in asthma compared to the healthy group across both cohorts (Figure 2I & 2J). The within-sample Shannon diversity of Anelloviridae was higher in asthma (p:0.024), while the Simpson index was significantly lower (p:0.008) in the MiSeq cohort, suggesting that healthy children had a few dominating species while asthmatic patients had a richer, more diverse and even distribution of Anelloviruses (Figure 2K & 2L); However, the difference was not significant in the HiSeq cohort (p>0.05).

### Divergent virome profiles linked with asthma and health

To further investigate the observed differentiation of the nasopharyngeal virome profiles in health and asthma we identified a set of 8 virome features in both discovery and validation cohorts (Figure 3A) and an additional 2 features in the latter (Figure 3B) that provided optimal hierarchical co-clustering of the healthy and asthma individuals; 70% (7/10) of healthy samples in the discovery cohort and 81% (9/11) of healthy samples in the validation cohort co-clustered (Figure 3A-3D). Then we focused on the sub-clusters that were obtained under the above configurations (Figure 3E & 3F). Based on the virome characteristics of these clusters (Figure S4 & S5) 3 virome profile groups (VPGs) were assigned: Prokaryotic VPG; (PVPG, n=32), contained samples with high prokaryotic richness and low/intermediate richness of eukaryotic viruses and Anelloviridae, Eukaryotic VPG (EVPG, n=12) included samples with high eukaryotic richness and low/intermediate Anelloviridae richness, and Anelloviridae VPG (AVPG, n=12) contained samples with high Anelloviridae richness (Figure 3E & 3F); We then investigated the relevance of the VPGs in asthma presence, severity, control and treatment: 81% of the healthy samples were grouped in PVPG compared to 42.8% of asthma samples (Figure 3G). We observed a gradient distribution of children with different levels of asthma control in the three VPGs (Figure 3H), while 57% and 24% of children with moderate and mild asthma, respectively, were grouped in AVPG, (Figure 3I). Treatments were evenly distributed among the three VPGs (Brown-Forsyth and Welch ANOVA tests p:0.66, p:0.63), suggesting that asthma treatment does not affect the composition of the VPGs. Overall, our data show a significant link between the nasopharyngeal virome, and the presence, control and severity of asthma in pre-school children.

**Figure 3:**
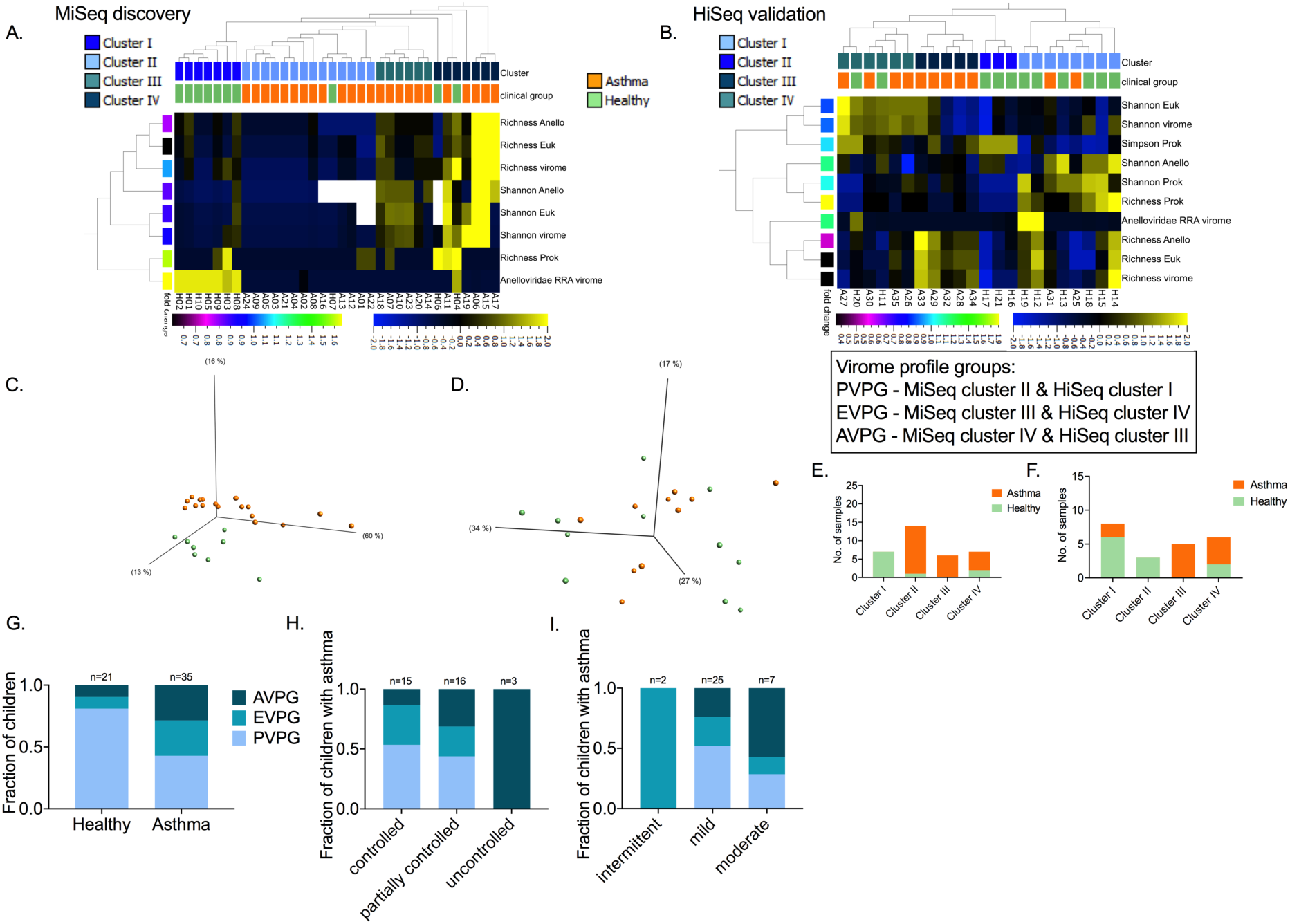
Virome profile groups identified in the MiSeq and HiSeq cohorts. Heat maps of the virome features that contributed to the co-clustering of healthy and asthma individuals in the (A) MiSeq cohort and (B) HiSeq cohorts; Eighteen virome features were filtered (feature reduction) based on their variance up to the level that provided the best health and asthma grouping; MiSeq: variance 0.11, projection score 0.18, and HiSeq: variance 0.04, projection score 0.13. The features that contributed to this separation are presented. All features were normalised to mean=0 and variance=1. Standardised heatmap color-coding; Yellow +2, Blue - 2. First row of sample annotation; Green: healthy, Orange: asthma. Second row of sample annotation: virome clusters; sample subgroupings based on the retained virome features. The cladogramms over the heat maps depict the agglomerative hierarchical clustering of individuals based on their virome characteristics (weighted linkage). Annotation of healthy and asthma individuals from the (C) MiSeq and (D) HiSeq cohorts in principal components. Stacked plots present the number and proportion of healthy and asthma individuals in each cluster identified in the (E) MiSeq and (F) HiSeq cohorts. Based on the virome properties of the samples in each MiSeq and HiSeq cluster, three overall virome profile groups were assigned: VPG1 (PVPG), VPG2 (EVPG) and VPG3 (AVPG). (G) Stacked plots of the fractions of healthy and asthma samples in the PVPG, EVPG and AVPG virome profile groups. (H) Stacked plots of the fractions of children with asthma in PVPG, EVPG and AVPG and different levels of asthma control: controlled, partially controlled and uncontrolled asthma. All cases of uncontrolled asthma were observed in AVPG along with 31.2% of partially controlled and 13.3% of controlled asthma. On the contrary, 43.7% of partially controlled and 53.3% of controlled asthma cases were observed in PVPG. (I) Stacked plots of the fractions of children with asthma in PVPG, EVPG and AVPG and different levels of asthma severity: intermittent, mild and moderate asthma.

### Dysbiotic structure of the asthmatic virome

To investigate the microbial community structure in health and asthma, we then studied the viral-viral and viral-bacterial co-occurrence. From our metagenomic sequencing, 355 bacterial metagenome assembled genomes (bMAGs) in the discovery and 522 in the validation cohorts have been identified. Their presence in health and asthma are depicted in Figure S6 and S7, respectively. We constructed two undirected microbial network meta-graphs representing an overview of microbial co-occurrences related to viruses in health and asthma (Pearson test p<0.05). The healthy meta-graph was composed of 328 nodes (MAGs) connected by 1180 edges (Figure 4A). The asthma meta-graph consisted of 326 MAGs with 1014 edges (Figure 4B). The normalised degree centrality, i.e. the number of interactions of each MAG, was significantly higher in the healthy meta-graph compared to the asthmatic (p<0.0001), suggesting reduced connectivity in asthma (Figure 4C). The Caudovirales, Anelloviridae and Picornaviridae viral families were the largest hubs in both the healthy and asthma meta-graphs, while taxa of the *Bacillales, Pseudomonadales, Enterobacterales*, and *Lactobacillales* order were the most highly paired bacteria with viruses (Figure S8A & S8B). Since prokaryotic virus richness was reduced in asthma, we investigated the co-occurring pairs of bacteriophages with bacteria (Figure 4D); in asthma, bacteriophages co-occurred with a significantly (p:0.0001) lower number of bacteria (Figure 4E) per bacterial order, compared to the healthy meta-graph, for most of the identified bacterial orders (Figure 4D). In the healthy meta-graph 71% (23/ 37) of virus-virus pairs involved taxa from different virus families (cross-family co-occurrence) (Figure S8C). In asthma, this was reduced to 40% (13/32) while 69% of them involved Enterovirus and Rhinovirus species A, B or C (Figure S8D). The above suggest that asthma is characterised by decreased connectivity (co-occurrence) within the virome-related microbiome, decreased number of bacteriophage-bacterial pairs, and increased cross-family co-occurrence of picornaviruses within the virome.

**Figure 4:**
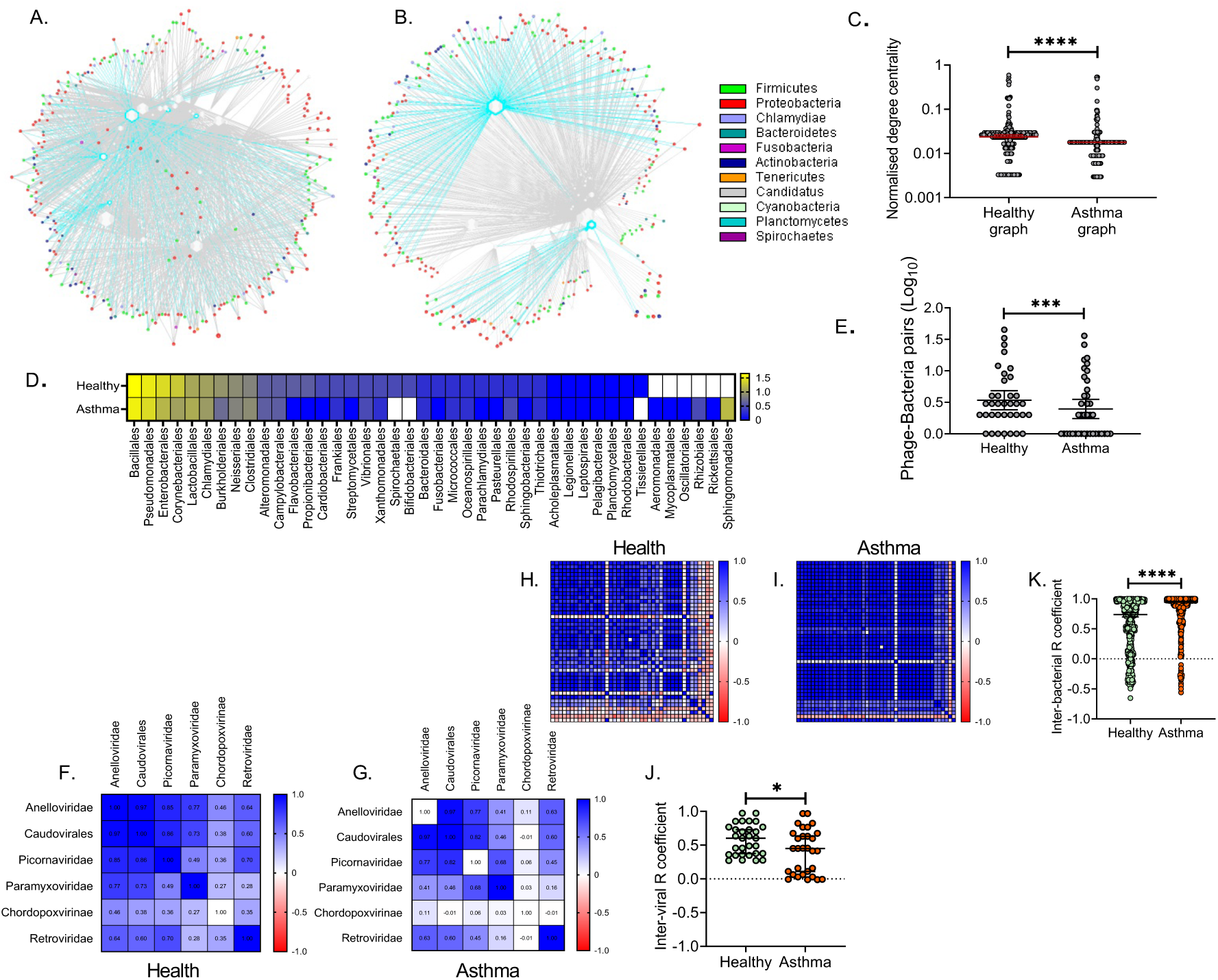
Viral-bacterial co-occurrence networks. (A) Healthy meta-graph, (B) Asthma meta-graph. Viruses are depicted as polygons in the centre of the graphs and bacteria as spheres in the periphery (virus concentric view). The size of each node is analogous to the normalised node degree centrality. Edges represent statistically significant correlations (Pearson test p<0.05) between viral and bacterial MAGs. Caudovirales nodes are highlighted with light blue as well as their edges. (C) Comparison of normalised degree centrality between the healthy and the asthma meta-graph; Statistical significance was assessed by the Mann-Whitney nonparametric t test. (D) Heatmap showing the log10 transformed number of phage-bacteria pairs in health and asthma and their distribution in bacterial orders (Yellow: high, Blue: low). (E) Comparison of the log10 transformed number of phage-bacteria pairs per bacterial order; Statistical significance was assessed by the Wilcoxon matched-pairs rank test. Color-coded pairwise correlation matrix amongst viruses identified in (F) health or (G) asthma in relation to their specific bacteria pairs (High: blue, Low: red). Color-coded pairwise correlation matrix amongst bacteria identified in (H) health or (I) asthma in relation to their specific virus pairs (High: blue, Low: red). (J) Comparison of the level of virus-virus correlation in respect to their bacteria pairs within the healthy and asthmatic meta-communities; Statistical significance was assessed by the Mann-Whitney nonparametric t test. (K) Comparison of the level of bacteria-bacteria correlation in respect to their virus pairs within the healthy and asthmatic meta-communities; Statistical significance was assessed by the Mann-Whitney nonparametric t test. Significance tests: * p<0.05, ** p<0.001, *** p<0.0001, **** p<0.00001. Scatter plots depict median values with 95%CIs.

To investigate inter-viral and inter-bacterial correlation with regard to the number and type of viral-bacterial co-occurrence, but also in relation to the fraction of interactions within the meta-graph, we expressed the number of specific virus-bacterial pairs as a fraction of the total number of edges (interactions) in each meta-graph and calculated the virus-virus (Figure 4F & 4G) and bacteria-bacteria (Figure 4H & 4I) pairwise R coefficients (Pearson correlations). In asthma, viruses correlated with each-other less positively than viruses in the healthy meta-graph (p: 0.031) (Figure 4J), based on their bacterial pairs. In contrast, the level of correlation amongst bacteria based on their virus-co-occurring pairs was significantly higher (p<0.0001) in asthma compared to the healthy meta-graph (Figure 4K). These observations suggest a less efficient structural organisation of the virome-related microbiome meta-community in asthma, characterised by reduced connectivity, fewer co-occurring phage-bacteria pairs, increased co-occurrence of viruses from different families—especially rhinoviruses—and increased correlation among bacteria. All the above are strong indications of virome dysbiosis in preschool children with asthma.

## Discussion

Despite the well-established effects of specific viral infections on the development, exacerbation and persistence of asthma^23-26^, the relationship between viral ecology of the airways and asthma remains poorly understood, and prokaryotic viruses in particular have received little attention. We show that during asymptomatic/infection-free periods, preschool children with asthma have a characteristically dysbiotic virome correlating with disease severity and control. The major component of dysbiosis is bacteriophage deficiency, while eukaryotic viral presence is increased, driven mainly by Anelloviruses and Picornaviruses. Networks of viral-viral and viral-bacterial co-occurrence were significantly loosened, suggesting that microbial communities may be less resilient in asthma. Viral dysbiosis therefore appears to be not only a key characteristic of asthma pathophysiology, but one that may be amenable to intervention^39-41^.

Viral exposures—and subsequent interactions with the host—are constant in the respiratory tract^42^. Furthermore, at least some viruses may persist^43^. Among these, prokaryotic viruses have been barely considered until now, despite their role in regulating bacterial abundance and promoting competition, stability and resilience^23,44-47^ within microbial ecosystems, which in turn may direct asthma development, activity and persistence^21,22,48,49^.

Bacteriophages are active regulators of bacterial populations^50-52^ and changes in phage composition or diversity have been linked to various diseases including Inflammatory Bowel disease, Parkinson’s disease and Type 1 Diabetes^53^. Moreover, bacteriophages can protect the epithelium from bacterial infections in a mucus-dependent manner, or can modulate the innate and adaptive arms providing non-host immunity^37,52,54-58^. In addition to reduced diversity and frequency of phage occurrence in asthmatics, the asthmatic metacommunity was characterised by a low number of phage-bacteria pairs, suggesting limited predator-prey ecological interactions^59-62^. Phage-bacteria dynamics is complex^63^ yet unexplored in the human airway and/or in relation to asthma. Microbial dysregulation through reduced phage presence is a novel concept in asthma pathophysiology. Changes in the bacteriome composition associated with risk of future asthma development and the loss of asthma control^16,21,48,64^ can potentially be linked to alterations in phage presence and/or dynamics. Our findings suggest that these scenarios are plausible, since virome profiles with reduced bacteriophages were associated with worsening of asthma control and severity.

The asthmatic virome was characterised by increased occurrence, richness and diversity of eukaryotic viruses. Since there is no reason to assume that exposure of children with asthma to viruses is governed by different epidemiological characteristics than healthy individuals, this could reflect differential levels of viral control through immune competence^65-67^; it is possible that asthmatics fail to clear eukaryotic viruses as efficiently as healthy children, allowing viruses to persist in the airways at low levels during asymptomatic periods. This may also explain a differential threshold for susceptibility to viral infection in asthma^68^. This may be further underpinned by the reduced evenness index of eukaryotic viruses in asthma, i.e. more than two species often contributing equally to the observed eukaryotic virome abundance, whereas the healthy eukaryotic virome was dominated by a single species even though more than two species could be detected.

Another association of the virome with the immune status can be drawn through Anelloviruses, which have been linked to functional immune competence^69-73^. We report that Anelloviruses are a stable component of the upper airway virome, as in other systems^69,74-76^, and in some cases they are observed with increased diversity and reduced evenness in asthma. The epidemiology of Anelloviruses is not entirely understood, nor are the factors that underlie their wide taxonomic diversity. It is possible that reduced immune competence may facilitate the colonisation of an individual’s airways with tight clusters of closely related Anelloviruses around founding clones (sometimes termed ‘quasispecies’). Moreover, in the asthmatic metacommunity, most of the viral-viral co-occurrence pairs involved Rhinovirus and Anellovirus species. Anelloviruses interact with the human host and hijack the host’s defence through direct suppression of NFκB, the regulation of TLR9 by viral-CpGs, and the regulation of IFN signalling pathways by virus-encoded micro-RNAs^75^. Notably, we have previously demonstrated that rhinoviruses are also encoded by genomes of extremely low CpG content which can influence TLR9-dependent stimulation^78-82^. This suggests that the immunomodulatory effect of Anelloviruses could act in favour for respiratory infections or condition the presence of sub-symptomatic low-virulence viruses to maintain a background level of chronic inflammation^70,71,83,84^. In support, we observed that the majority of asthma donors had an Anellovirus-rich (AVPG) or eukaryotic-rich (EVPG) virome and that compromised asthma control was associated with these virome profiles.

The reduced connectivity within the asthmatic virome-related metacommunity suggests a reduction of species co-occurrence and/or increased interpersonal variability. Increased microbiome heterogeneity has been suggested to be a hallmark of dysbiotic microbiomes^14,85^, and our virome data strongly point in this direction. Interestingly, virus-based interbacterial correlations were greater in the asthmatic metacommunity, and lacked negative correlations. This suggests reduced modularity among microbes co-occurring with viruses, and thus a reduced ability to compartmentalise virus-induced microbiome perturbations in asthma, such as during the addition or elimination of a virus from the system^86^. This type of microbiome dysregulation could lead to reduced stability, resilience and shifts in the microbial composition upon perturbations^44,87,88^, for example during respiratory infections and/or virus-induced asthma exacerbations. Notably, differential pathogenic bacteria-to-bacteria correlations have also been identified in tonsils of atopic individuals^89^.

In this study, a number of choices had to be made regarding the design, analysis and interpretation of the metagenomic data. First, metagenomic sequencing allows the identification of viral and microbial genomes or genome ‘traces’, without however concluding presence of infectious pathogens^90-92^. Viral enrichment through gradient ultra-centrifugation can bias the sequencing output as it depends on previous knowledge of the physical properties of the virions or virus like particles^93,94^. To tackle the above, we followed a sample-processing strategy filtering out ‘naked’ nucleic acids while retaining encapsulated viral sequences of DNA and RNA genomes^95,96^, in order to be as close as possible in the identification of virus-like particles or viruses. Second, the sequencing strategy, as demonstrated, can have a significant effect in the identification of viral genomes and the subsequent meta-analysis. Sequencing too deep can blur a differentiating signal while sequencing at a low depth reduces diversity. To address this, we have used two different sequencing strategies and asked whether we can observe the same or similar differences between health and asthma. This was also expanded to different geo-locations including potentially variable virus exposure characteristics. Third, we chose to focus at the ecological (community) level. The ecological indices used are either incidence- or abundance-based. Viruses do not follow uniform replication/infection kinetics and have different hosts, i.e. eukaryotic versus prokaryotic viruses. In contrast to bacteriome studies where the relative abundance of each species implies some level of time-dependent stable presence in the system, the same cannot be assumed for viruses. Even, at a longitudinal setting, unequal time-intervals would be needed to study stable viral presence. Thus, our data provide a snapshot view of virus presence in the nasopharynx. Finally, the cohorts comprised of preschool-age children, an age during which the diagnosis of asthma is challenging and prognosis uncertain. Nevertheless, asthma persists in a considerable proportion of such children providing the opportunity for evaluating further outcomes. As with other microbiome niches, the respiratory virome is expected to change with time; whether the observed pathophysiology is age-specific or not remains to be elucidated. This is also the case for different asthma phenotypes and severity levels. Nevertheless, the PreDicta cohort was shown to be representative of preschool asthma^27^. Finally, the identified viromes and bacteriomes refer to the specific tissue niche of the nasopharynx and further investigation is needed to study viral presence along the respiratory tract.

Overall, we report novel findings in the virome of children with asthma with potential for intervention. Reduced phage presence and/or diversity might lead to dysregulated bacterial colonisation through the absence of appropriate ecological interactions compromising the system’s resilience to environmental^6,97-99^ or immune-driven perturbations like asthma exacerbations^49,100^. In the same direction, increased co-occurrence of eukaryotic viruses might increase the chance of a symptomatic infection or sustained inflammation. These events could induce stochastic changes^85^ in the respiratory metacommunity triggering a transition to compositional states^16,21,49,64,101^ with increased susceptibility for viral and bacterial infections, which in turn further compromise the system’s stability and resilience. The temporal trajectories of these complex dynamics and their association with immune competence and disease persistence at different ages, are currently explored by the CURE consortium^102^.

## Acknowledgements

We thank Dr Anna Sobanska for her valuable contribution in patients’ recruitment and care at Lodz center, Poland.

## Author contributions

Conceptualization, SM, NP; Methodology, SM, BC, AT, DR, NP; Software, SM, BC, AT, AGS; Investigation, SM, DR, NP.; Resources, VX, MK, CB, SNF, TJ, TV; Writing – Original Draft, SM, DR, NP; Writing – Review & Editing; SM, BC, AT, AGS, VX, MK, CB, SNF, TJ, TV, EA, DR, NP; Visualization, SM, BC.; Supervision, DR, NP.

## Methods

### Cohort description

Donors were recruited as part of the Predicta cohort^103^. The PreDicta paediatric cohort was a two-year prospective multicentre study (five European regions) under the EU FP7 program. The cohort was designed to prospectively evaluate wheeze/asthma persistence in pre-schoolers associated with viral/microbial exposures and immunological responses^103^. Children with an asthma diagnosis^104^ of mild to moderate severity according to GINA^105^ within the last two years were recruited. Donors did not present an episode of an URT and/or asthma exacerbation during the preceding four weeks prior to inclusion and baseline evaluation. Since all donors were retrospectively monitored for two years, detailed knowledge of upper (URTIs) and lower (LRTIs) respiratory tract infection events following baseline evaluation. In this study we sought to characterise the nasopharyngeal metagenome of preschool children at a baseline, non-infectious and non-exacerbation state, with emphasis on the viral component (virome). Thus, the timing of sample collection was at least one month prior to and one month following a reported respiratory tract infection. Nasopharyngeal samples were included from two groups of donors: a discovery group from southern Europe (Athens, Greece), and a validation group comprising individuals from Lodz (Poland) and Turku (Finland) (Central and North Europe). Except for geographical origin, clinical characteristics were similar among donors from the two groups (Supplementary table 1). The discovery cohort included 10 healthy donors and 24 asthma patients. The validation cohort included 11 asthmatic patients and 11 healthy donors (Supplementary table 1).

**Supplementary table 1:**
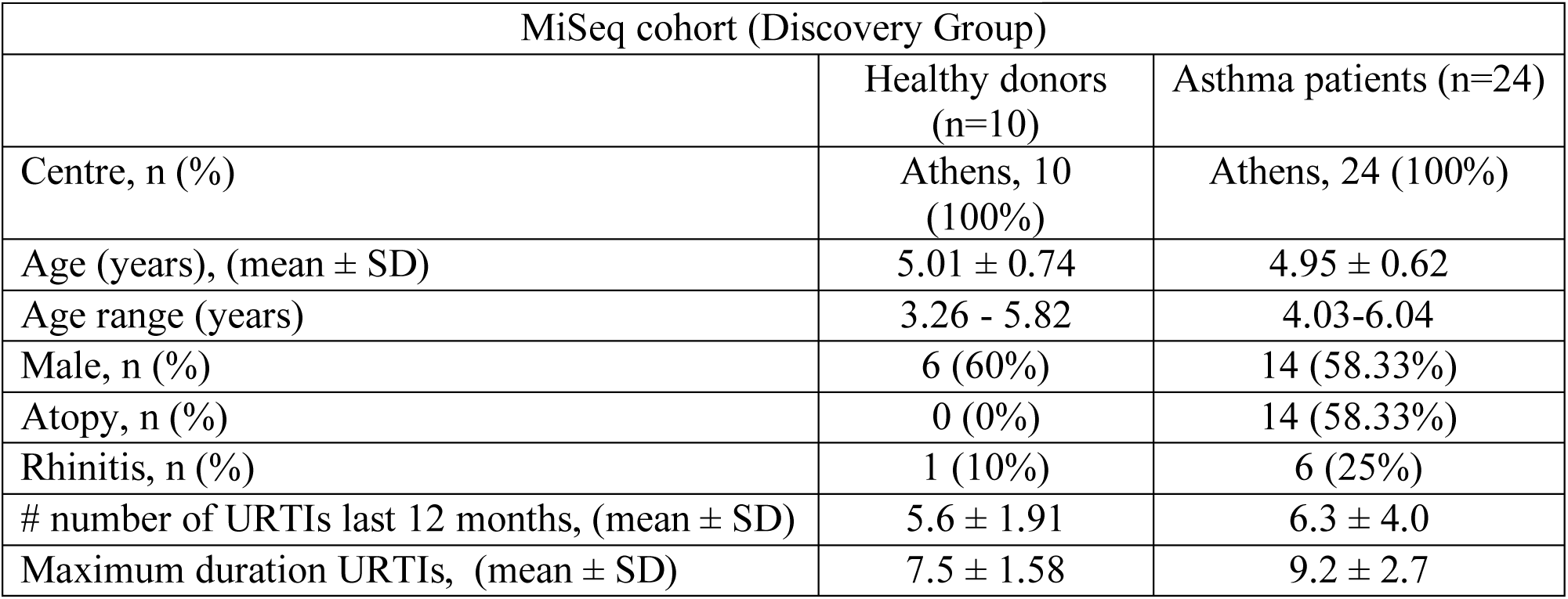
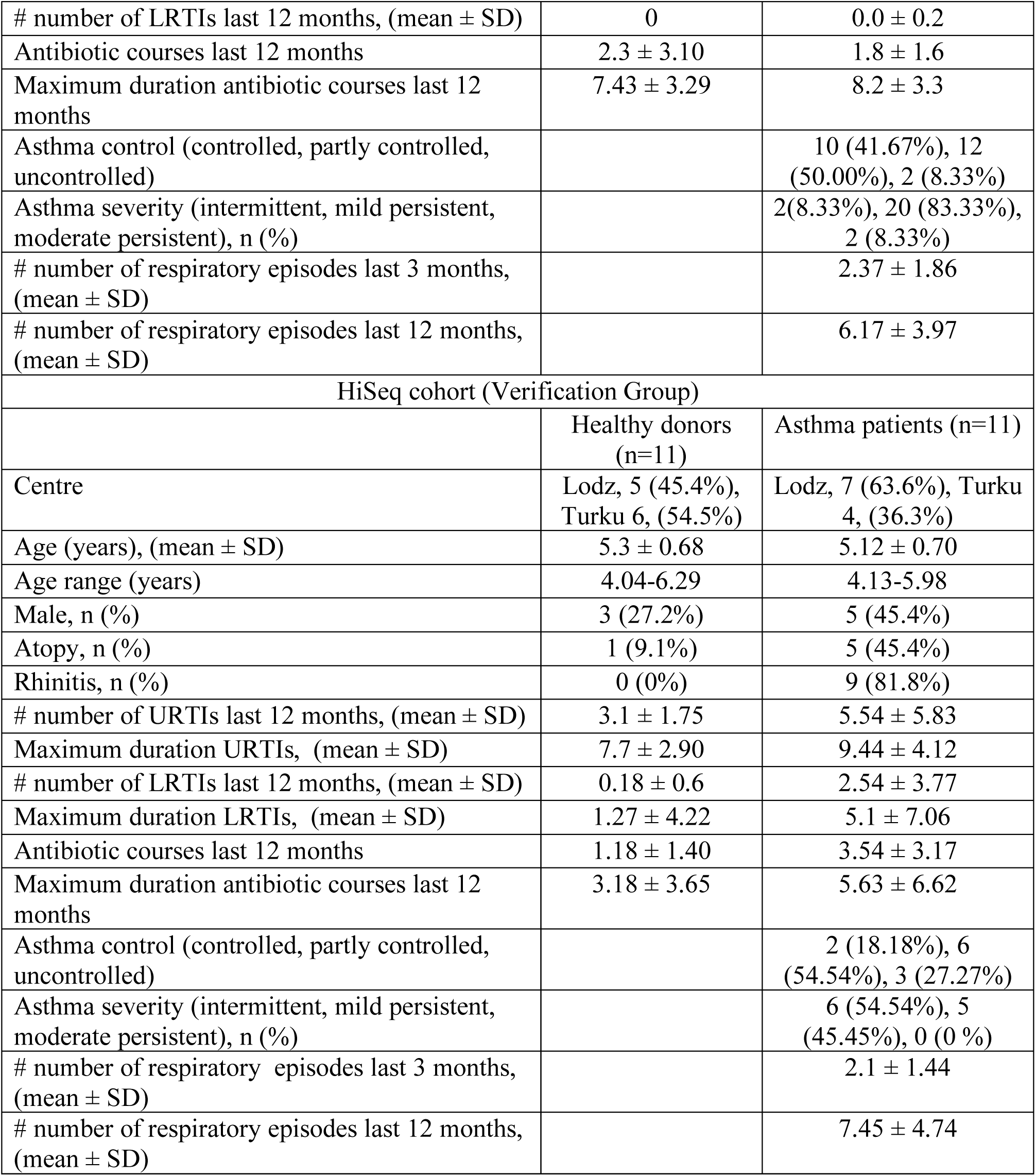
Characteristics of study participants.

### Sample processing

Nasopharyngeal samples were obtained using flocked nasopharyngeal swabs (ESwab™ collection and transport system, COPAN)^106^. Specimens were collected using the same standard operational procedure (SOP) across the recruitment centres. The nasopharyngeal specimen was eluted in 1000 µl of transport medium (Amies buffer), split into 200 µl aliquots and stored at −80°C. One aliquot was sent to the University of Manchester for metagenomic analysis. Samples obtained from asthma patients and healthy participants were analysed in parallel to reduce bias. Samples were slowly defrosted and centrifuged under chill conditions at 2500g for 5 minutes to spin down cells and debris^107^. Caesium chloride purification was not used because it can lead to bias towards specific virion like particles (VLPs)^93,94^. To enrich for encapsulated nucleic acids, supernatants were split into four aliquots (50 µl) and treated with a mixture containing 5 units of Turbo DNAse (Ambion-ThermoFisher Scientific) and 2.5 µl RNAse inhibitor (RNaseOUT™, Invitrogen-ThermoFisher Scientific) in DNAse reaction buffer (Ambion-ThermoFisher Scientific) and incubated for 45 minutes at 37°C^108^. Ten microliters of DNAse inactivation reagent were added to each aliquot and incubated for 5 minutes at room temperature. Samples were centrifuged at 10,000g for 1.5 minutes and supernatants were pooled back together per sample (total of 180 µl) and used for subsequent nucleic acid isolation. All samples were processed in a class 2 biosafety cabinet to reduce environmental contamination. A negative control sample (sterile filtered water) was introduced to each experiment, followed the whole protocol and the DNA concentration was measured before library preparation using a Qubit™ fluorometer (ThermoFisher Scientific). Negative controls did not produce measurable readings (estimated concentration <10 pg/µL).

### Metagenomic nucleic acid extraction

Nucleic acids were extracted using the AllPrep DNA/RNA Micro kit (Qiagen) for simultaneous purification of minute amounts of DNA and RNA from the same sample without making use of the carrier RNA. An additional on-column DNAse treatment step (RNAse free DNAse set, Qiagen) was added to remove possible traces of remaining DNA co-extracted with the RNA. The quality of DNA and RNA was evaluated in a NanoDropTM1000 spectrophotometer (ThermoFisher Scientific) and the yield in the Qubit™ fluorometer (ThermoFisher Scientific) using dsDNA and ssRNA high sensitivity assays (ThermoFisher Scientific). Synthesis of the first strand of cDNA was carried out using the SuperScript™ II reverse transcriptase system and random primers (ThermoFisher Scientific) with default conditions. The second strand of the cDNA was synthesised using Klenow fragment polymerase (New England Biolabs). Ten microliters of cDNA were incubated for 2 min at 95°C and chilled on ice for 2 min before addition of 5 units of Klenow fragment and incubation at 37°C for 1 h. Enzyme inactivation was performed at 75°C for 10 min. The resulting product was a double-stranded copy of the initial single-stranded cDNA. To amplify the ds-cDNA and genomic DNA, a whole genome amplification strategy was carried out using multiple displacement amplification^109,110^, TruePrime™ WGA kit (SYGNIS)^111-113^. Samples were incubated for 3 hours at 30°C followed by polymerase inactivation at 65°C for 10 minutes. DNA amplification yield was confirmed by fluorometric measurement as described before. The whole process negative control samples did not produce any measurable readings (estimated concentration<10 pg/µL). Overall, each swab specimen produced two amplified dsDNA samples, one derived from the initial isolated DNA (metagenomic DNA sequences) and a second from the initial isolated RNA (metagenomic RNA sequences).

### Metagenomic high throughput sequencing

An aliquot of 10 µl (Normalisation: 1 ng of DNA per reaction) from each sample was transferred to the Genomic Technologies Core Facility (GTCF) in the University of Manchester. Illumina sequencing libraries were generated using ‘on-bead’ tagmentation chemistry with the Nextera DNA Flex Library Prep Kit (Illumina, Inc.) according to the manufacturer’s protocol. For the MiSeq cohort, DNA libraries were pair-end sequenced using the MiSeq high throughput sequencing instrument (Illumina) in 2×75 base pair (bp) runs. For the HiSeq cohort DNA libraries were pair-end sequenced using the HiSeq4000 high throughput sequencing platform (Illumina) in 2×150 bp. Each flow cell contained libraries from both asthma patients and healthy children to avoid sequencing bias. The output data was de-multiplexed and BCL-to-FASTQ conversion was performed using Illumina’s bcl2fastq software, version 2.17.1.14.

### De novo assembly, microbial annotation and taxonomy assignment

We used customised computational strategies to achieve *de novo* assembly of contigs from short read sequences: In the MiSeq cohort (MiSeq, low read output), reads with alignments to host sequences (hg38) were discarded (BBMap 36.92) and remaining reads were trimmed of adapter sequences and low-quality regions (BBDuk 36.92). Filtered paired and orphan reads were pooled across all individuals and co-assembled (MetaSPAdes 3.9.1; k-mers 17,21,25,31,43,55,67; doi:10.1101/gr.213959.116) before being queried against the NCBI NR protein database dated 2017-01 (Diamond 0.8.34; doi:10.1038/nmeth.3176). Taxonomic ranks were assigned to each contig from protein homology search results using the weighted lowest common ancestor algorithm (wLCA) https://arxiv.org/abs/1511.08753 as implemented in MEGAN 6 (doi:10.1101/gr.5969107) using a 50% match weighting. Contigs assigned known non-microbial (namely host) taxa were discarded. Finally, reads from individuals were mapped to combined meta-assembly contigs (BBMap 36.92; seed k=7) representative of i) entire microbiome and ii) viruses, enabling inference of relative taxonomic abundances for each individual. More than 70% (78.92%) of contigs were classified, mapping to 380 distinct microbial taxa including 25 viral metagenomic assembled genomes (vMAGs). A total of 214,533,315 sequencing reads from 34 complete metagenomic specimens (Mean log_10_: 5.785±SEM 0.154) were aligned back into MAGs. 5,351,667 sequencing reads aligned to 25 vMAGs (Mean log10: 2.156±SEM 0.434).

In the HiSeq cohort (HiSeq, high read output), BBTools (38.23) were used to decontaminate libraries from human sequences and remove artefacts (BBDuk and BBMap, sliding window based quality filtering at Q10). Filtered paired and orphan reads were co-assembled and individual-based assembled. For co-assembly, sequencing reads from DNA & RNA-derived samples were assembled with MetaSpades (3.12.0) at k-mers k 21, 33, 55, 77, 99, 127. Megahit (1.1.3) was used for individual-based assembly with default K-mers (k 21, 29, 39, 59, 79, 99, 119, 141). MetaQUAST was used to merge co-assembled and individual-based assemblies. LCA was performed in DIAMOND 0.9.22 (NR database and taxon mapping downloaded 2018-09-13). Sequencing reads remapped (BBMap, default settings with scafstats output) to all MetSPAdes metagenomic assembled genomes. Overall, we produced a total of 34,031 classified contigs, organised in 576 MAGs, including 54 viral MAGs (vMAGs). About 700 million sequencing reads (712,176,554) from 22 complete metagenomic specimens aligned onto the microbiome MAGs (Mean log_10_: 6.684±SEM 0.113). Of those, 121,694,232 sequencing reads aligned to vMAGs (Mean log_10_: 4.782±SEM 0.299). A summary of the complete microbiome MAGs identified in healthy and asthmatic children can be seen in Figure S6 and Figure S7, respectively. Viral genomes were identified within the ENA sequence viral database using EBI’s BLASTN REST API^114^ and 7mer MASH distances were generated with Sourmash^115^ (Figure S1G).

### Estimation of vMAG ecological indexes

To avoid confounding effects of sample size and underestimation of true species richness, we have performed sample-based randomisation and rarefaction with extrapolation. We have evaluated the asymptotic richness estimator at all levels of taxonomic rank accumulation (rarefaction) up to the size of the reference sample: MiSeq cohort (healthy: *n*= 10, asthma: *n*= 24), HiSeq cohort (healthy: *n*= 11, asthma: *n*= 11) using 100 randomizations. The means (and conditional standard deviations) among resamples for each level of accumulation were reported. Extrapolation was performed by a factor of 1 (EstimateS, version 9.1.0)^116^. For comparison of the sample-based rarefaction curves (richness divergence between clinical groups), non-overlap of 95% confidence intervals constructed from the unconditional variance estimators is used as a simple but conservative criterion of statistical difference^116^. To define within-donor (individual-based) vMAG alpha diversity we have used the number of vMAGs observed per analysed specimen, the abundance-based Shannon diversity and the abundance-based Simpson evenness^117^. For abundance-based ecological indexes, vMAG abundance was computed using the relative read abundance; the number of sequencing reads per vMAG was normalised against the total number of sequencing reads observed per taxonomical group or any other type of grouping, e.g. eukaryotic and prokaryotic viruses. We computed the between-samples diversity for each assemblage using incidence-based (Jaccard and Sorensen) measures of relative compositional similarity (EstimateS, version 9.1.0)^116^. For incidence-based similarity, the Jaccard index compares the number of shared species to the total number of species in the combined samples of each assemblage (global view), while the Sorensen index compares the number of shared species to the mean number of species in a single assemblage (local view)^118^.

### Species co-occurrence networks

We removed singletons and doubletons per each clinical group and sequencing strategy and retained MAGs with at least 3 occurrence events. We calculated the pairwise R coefficients of the MAGs based on their relative sequencing read abundance within the metagenome. The pairwise comparison matrices were transformed into interaction graphs: each node represented a microbial lineage in the LCA (MAG) and an edge represented a positive or negative correlation between two lineages (MAGs). To dichotomize the presence or absence of a co-occurrence link, only significant (Pearson test p<0.05) pairwise correlations involving a vMAG were selected; these included 40 co-occurring pairs in the healthy and 36 in the asthma MiSeq cohort, and 1765 pairs in the healthy and 1412 in the asthma HiSeq cohort. All graphs were visualised using Network Analysis, Visualization, & Graphing TORonto (NAViGaTOR)^119^. Networks were organised in order to separate different components using the GRIP algorithm or the concentric view option. Node and edge descriptives were analysed using NAViGaTOR.

### Statistics

Continuous variables were tested for normal distribution using the Shapiro-Wilk test. Pairwise quantitative differences between two groups were tested using two-tailed t test and Mann Whitney (Wilcoxon rank sum test) for variables not following a normal distribution. F test was used to test for equal variance. Pairwise comparisons between three or more groups was performed with the Brown-Forsythe and Welch ANOVA tests corrected for multiple comparisons by controlling the False Discovery Rate (FDR) for normally distributed variables and the Kruskal-Wallis test corrected for multiple comparisons (Dunn’s test) for variables not following a normal distribution. Principal component analysis, hierarchical agglomerative clustering (average and weighted linkage), K-means clustering and feature reduction based on variance was performed on normalised variables with mean of 0 and variance of 1 (Qlucore Omics explorer, version 3.4); We used principal component analysis and exploratory visualization to find a representation from which we can extract interpretable and potentially interesting information, following the evolution of the projection score in real time during variance filtering (Qlucore Omics explorer, version 3.4). The Pearson test was used to test for pairwise correlations among continuous variables. Statistical tests and graphical visualisations were produced in GraphPad prism 8.02.

---

**Supplementary figure 1:**
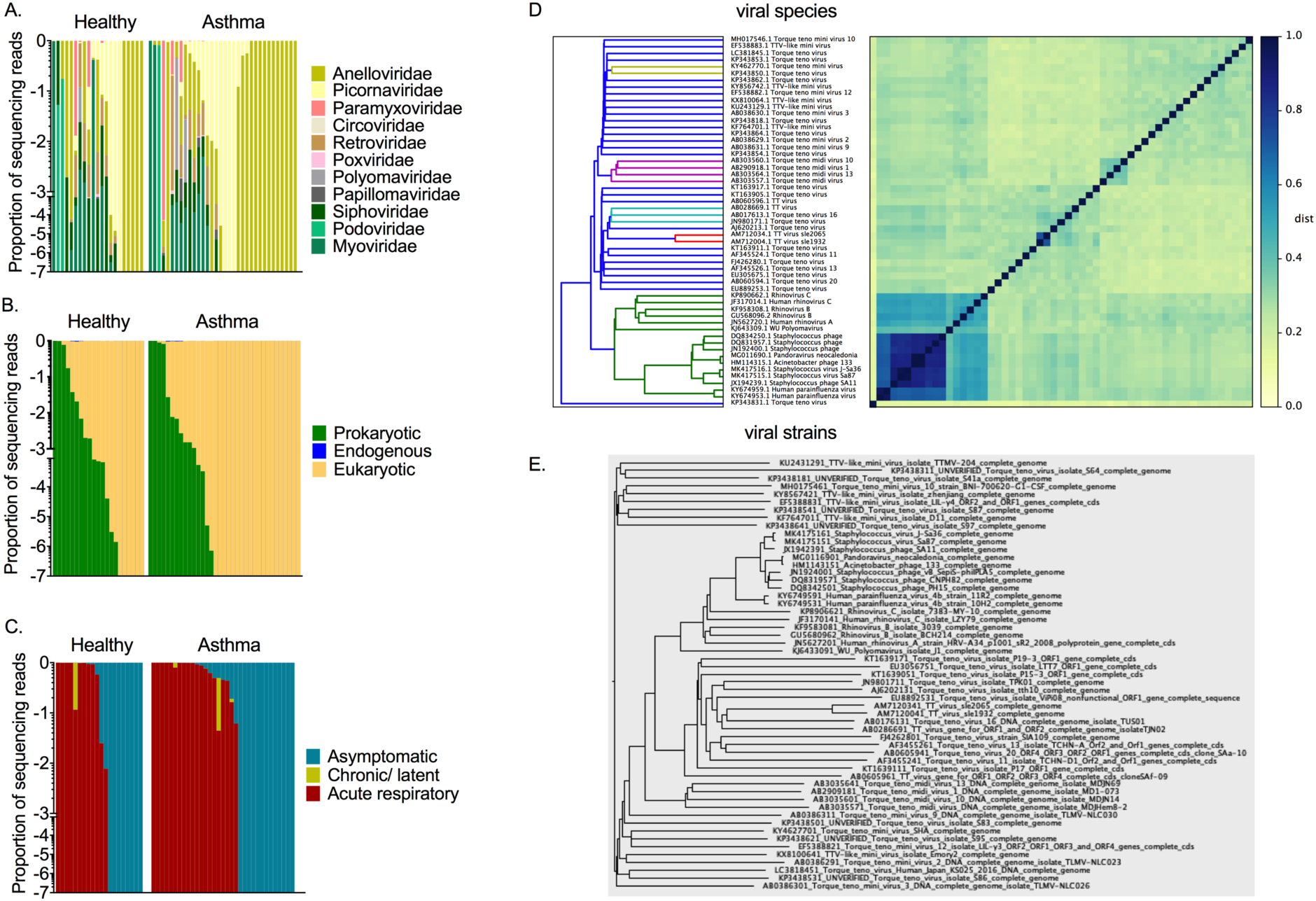
Virome description. Virus abundance plots per healthy and asthmatic individuals. Viruses are organised based on different properties: (A) taxonomic family, (B) host-type, and (C) type of pathogenicity. (D) Clustered Heatmap and dendrogram of 7mer MASH distances between vMAG representative genomes at the species, and (E) strain level. Nonredundant representative genomes for each vMAG were identified within the ENA Sequence Viral database using nucleotide BLAST.

**Supplementary figure 2:**
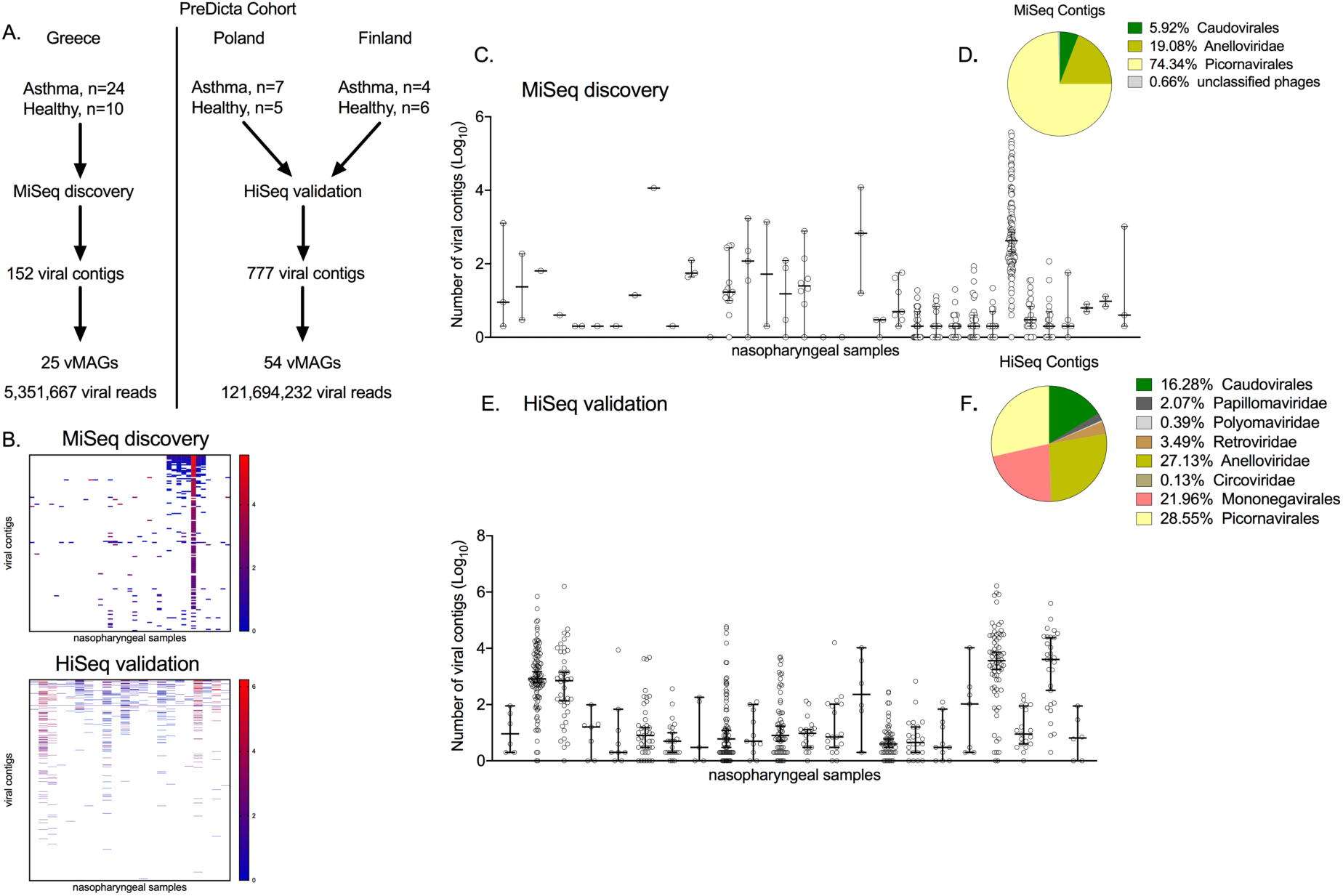
Viral metagenome assembled genomes identified in the study. (A) Study design and metagenomic sequencing output. (B) Heat maps of the number of sequencing reads aligned into viral contigs identified in the nasopharyngeal samples of children screened with MiSeq and HiSeq; Columns: samples, rows: viral contigs; High: red, low: blue. Scatter plots of the number of different viral contigs identified in each sample sequenced in (C) MiSeq, and (E) HiSeq; Median values with 95%CIs are depicted. Pie charts of the fraction of viral contigs classified at different taxonomy levels in (D) MiSeq, and (F) HiSeq.

**Supplementary figure 3:**
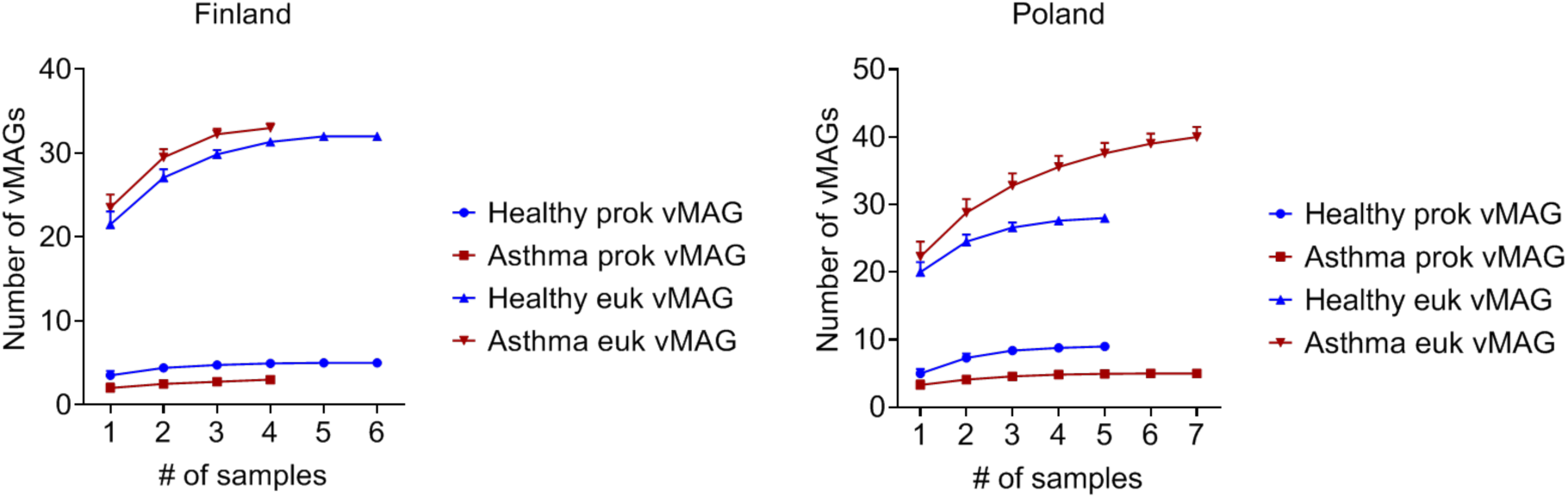
Estimation of viral richness in donors from Finland and Poland. The number of vMAGs was estimated using sample-based rarefaction curves without replacement (100 iterations) with increasing number of subsamplings. The plots depict the estimated number of vMAGs for the prokaryotic and eukaryotic viruses in healthy and asthma donors recruited in Turku (Finland) and Lodz (Poland). The mean and standard deviation (SD) is depicted for each data point. Dark red: Asthma, Blue: healthy.

**Supplementary figure 4:**
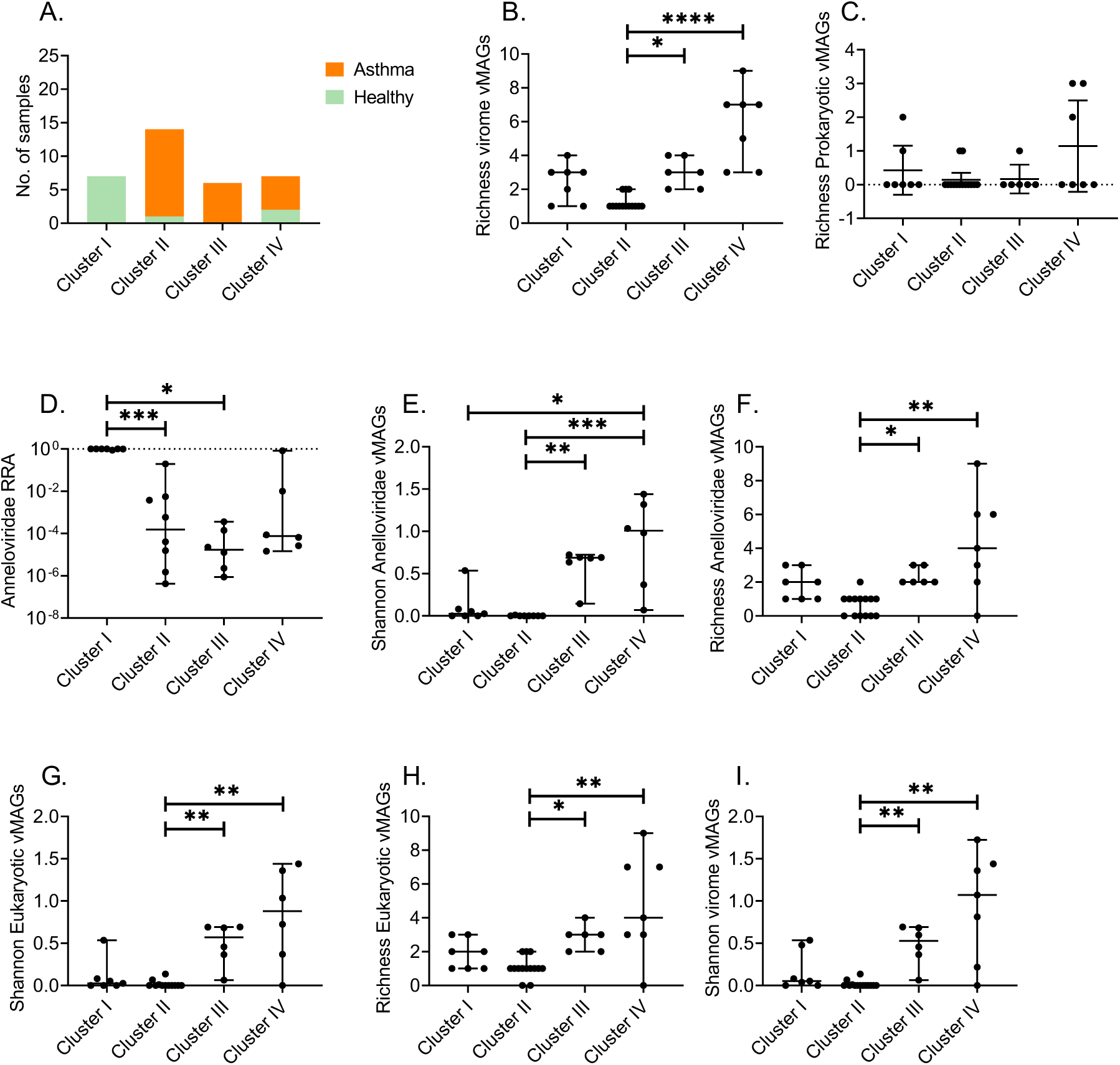
Clusters of donors with different virome properties identified in the MiSeq cohort. Separation of healthy and asthma donors was based on eight virome features (Figure 4A). Based on these features 4 subgroups of donors were identified. The fraction of healthy and asthma samples in each cluster is depicted in (A). The quantitative differences between the samples of each cluster for the 8 virome features are depicted: (B) Richness of the total virome, (C) Richness of the prokaryotic virome (bacteriophages), (D) Relative read abundance of the Anelloviridae family, (E) Shannon abundance-based diversity of Anelloviruses, (F) Richness of Anelloviruses, (G) Shannon diversity of the eukaryotic virome, (H) Richness of the eukaryotic virome, and (I) Shannon diversity of the virome. Kruskal-Wallis test corrected for multiple comparisons (Dunn’s test); adjusted p values are reported: Significance tests: * p<0.05, ** p<0.001, *** p<0.0001, **** p<0.00001. Scatter plots depict median values with 95%CIs.

**Supplementary figure 5:**
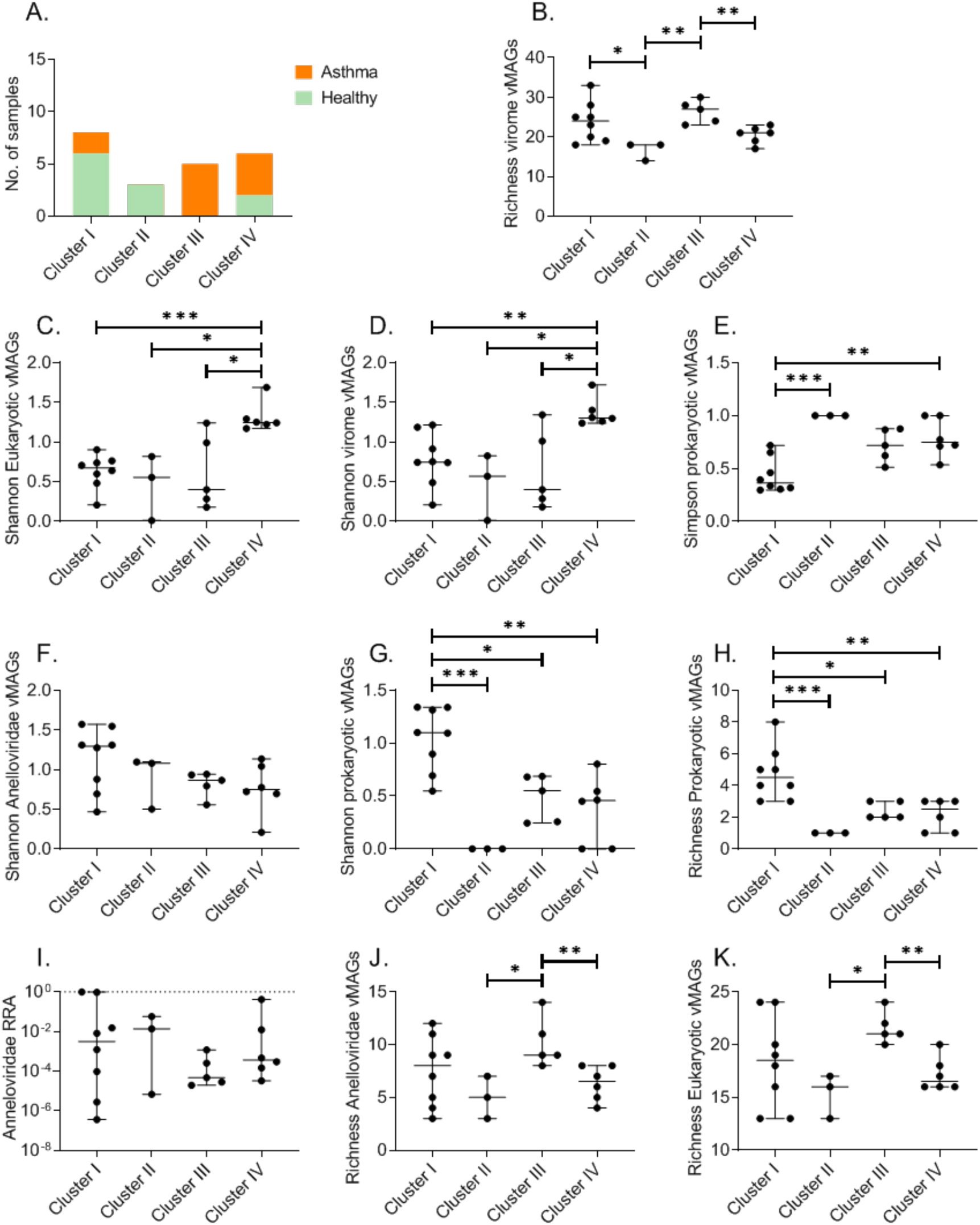
Clusters of donors with different virome properties identified in the HiSeq cohort. Separation of healthy and asthma donors was based on ten virome features. Based on these features 4 subgroups of donors were identified. The fraction of healthy and asthma samples in each cluster is depicted in (A). The quantitative differences between the samples of each cluster for the 10 virome features are depicted: (B) Richness of the total virome, (C) Shannon abundance-based diversity of the eukaryotic virome, (D) Shannon diversity of the virome, (E) Simpson abundance-based evenness of the prokaryotic virome, (F) Shannon diversity of Anelloviruses, (G) Shannon diversity of the prokaryotic virome, (H) Richness of the prokaryotic virome, (I) Relative read abundance of Anelloviruses, (J) Richness of Anelloviruses, and (K) Richness of the eukaryotic virome. (B, C, D, J, K) Brown-Forsythe and Welch ANOVA test corrected for multiple comparisons by controlling the False Discovery Rate (FDR). (E, F, G, H, I) Kruskal-Wallis test corrected for multiple comparisons (Dunn’s test). Adjusted p values are reported: Significance tests: * p<0.05, ** p<0.001, *** p<0.0001, **** p<0.00001. Scatter plots depict median values with 95%CIs.

**Supplementary figure 6:**
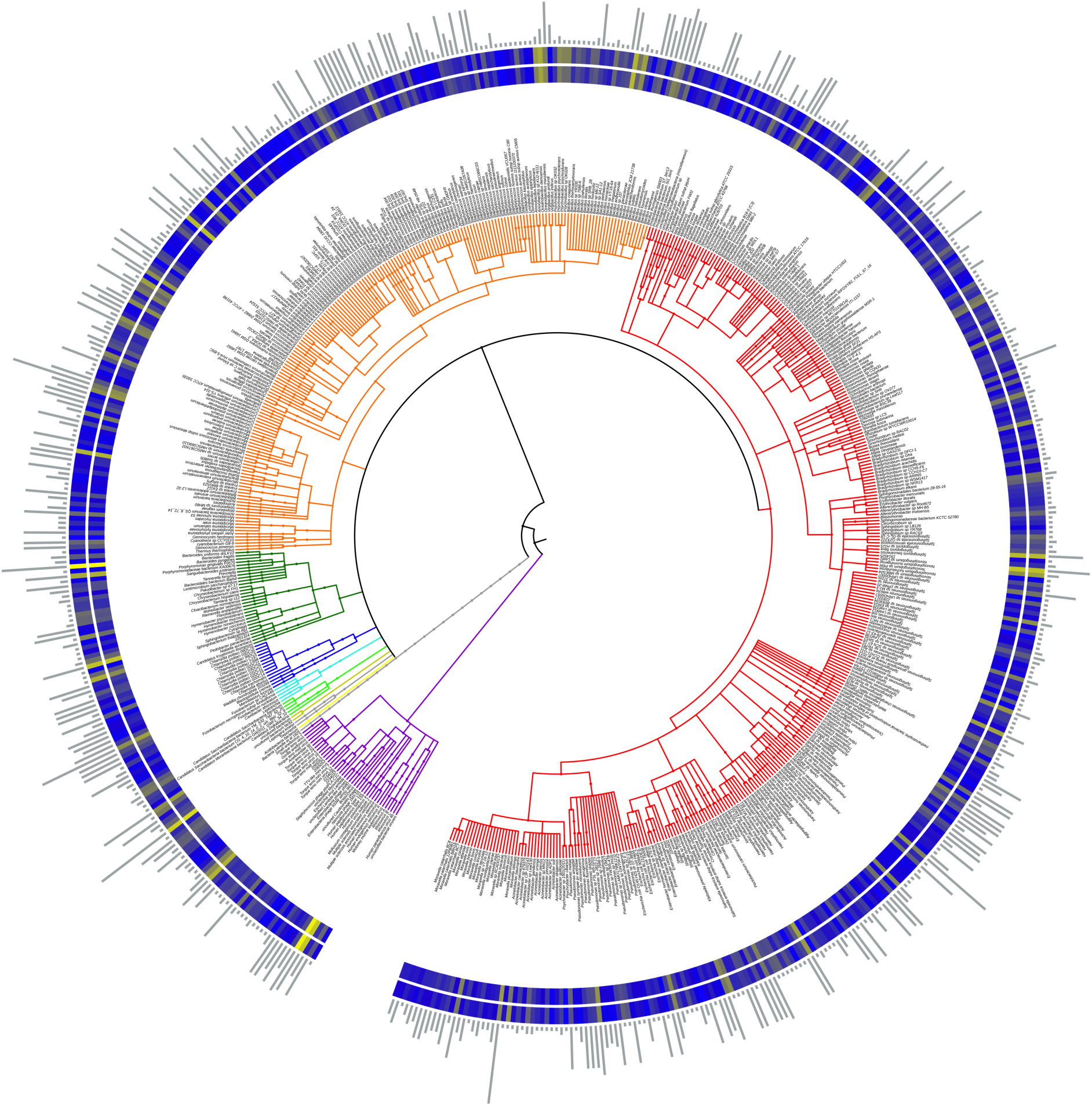
Respiratory metagenome assembled genomes (MAGs) of healthy children. MAGs are organised into hierarchical clusters based on their taxonomic similarity (taxonomic lineage up to species level). A total of 813 (95%CI: 141-241) microbial taxa were identified in healthy children; 757 out of 813 taxa belonged to bacteria and 56 to viruses. We observed 236 (29%) microbial taxa, including 14 viral taxa, present exclusively in the healthy group. About 369,655,276 (mean log10: 6.610±0.972, n=21) sequencing reads aligned to microbial MAGs isolated from healthy samples. The mean and total relative read abundance of each MAG are depicted as colour gradient concentric circles around the cladogramm (log10 transformed), respectively. The frequency of occurrence for each lineage (up to species level) is visualised as bar plots in the outer ring of the circular cladogramm. Cladogramm colouring: Purple; Viruses, Red; Proteobacteria, Orange; Firmicutes, Green; Bacteroidetes, Dark Blue; Chlamydiae, Light Blue; Fusobacteria, Light Green; Candidatus bacteria, Gold; Unclassified bacteria, Light Yellow; Acidobacteria

**Supplementary figure 7:**
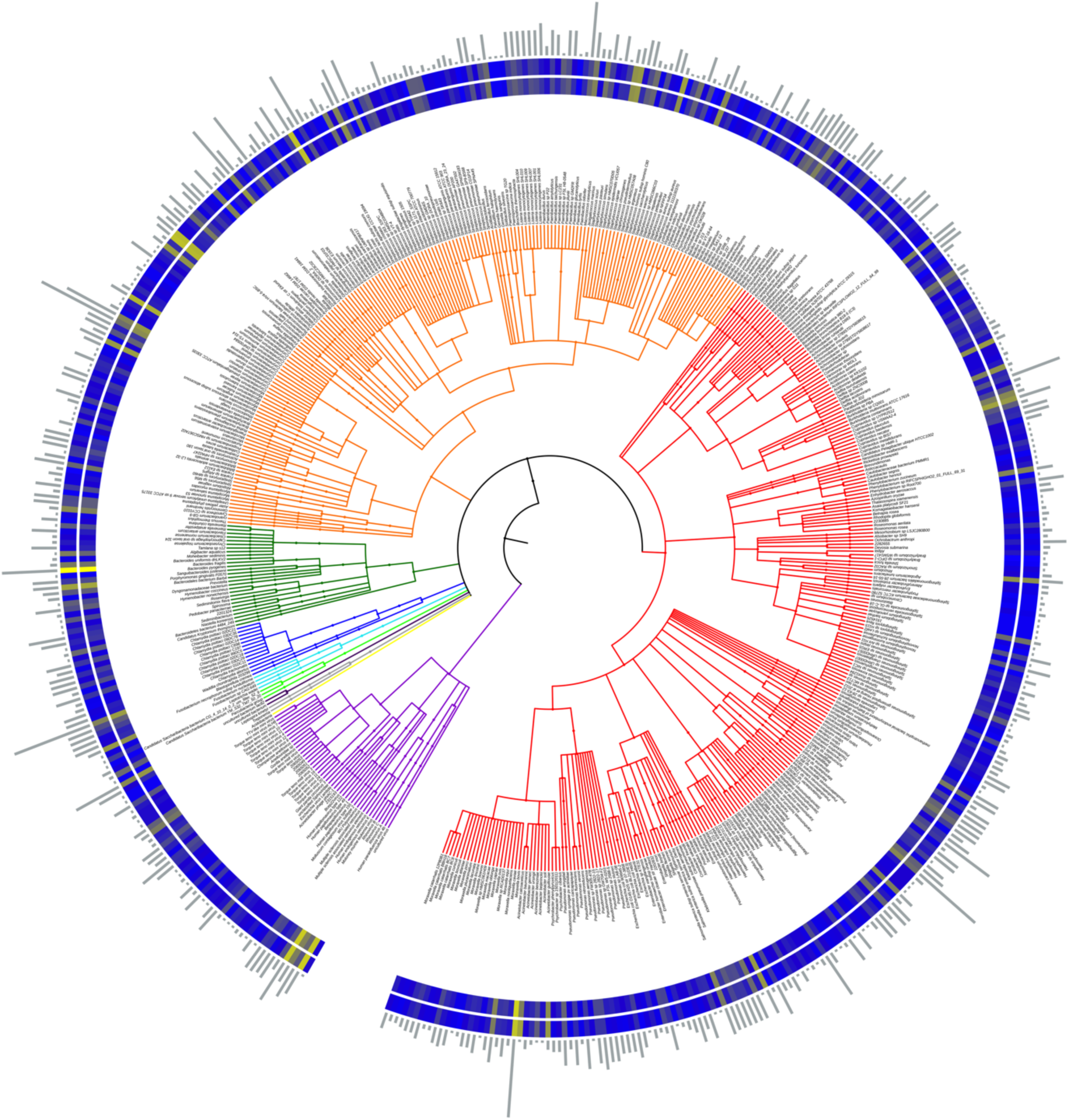
Respiratory metagenome assembled genomes (MAGs) in children with asthma. MAGs are organised into hierarchical clusters based on their taxonomic similarity (taxonomic lineage up to species level). A total of 722 (95%CI: 102-177) microbial taxa were identified in asthma; 662 out of 722 taxa belonged to bacteria and 60 taxa to viruses. We observed 144 (20%) taxa, including 18 viral taxa, unique for asthma patients. 490,164,409 (mean log10: 6.273±1.164, n=35) sequencing reads aligned to MAGs isolated from asthma samples. The mean and total relative read abundance of each MAG is viewed as colour gradient concentric circles around the cladogramm (log10 transformed), respectively. The frequency of occurrence for each lineage (up to species level) is visualised as bar plots in the outer ring of the circular cladogramm. Purple; Viruses, Red; Proteobacteria, Orange; Firmicutes, Green; Bacteroidetes, Dark Blue; Chlamydiae, Light Blue; Fusobacteria, Light Green; Candidatus bacteria, Gold; Unclassified bacteria, Light Yellow; Acidobacteria.

**Supplementary figure 8:**
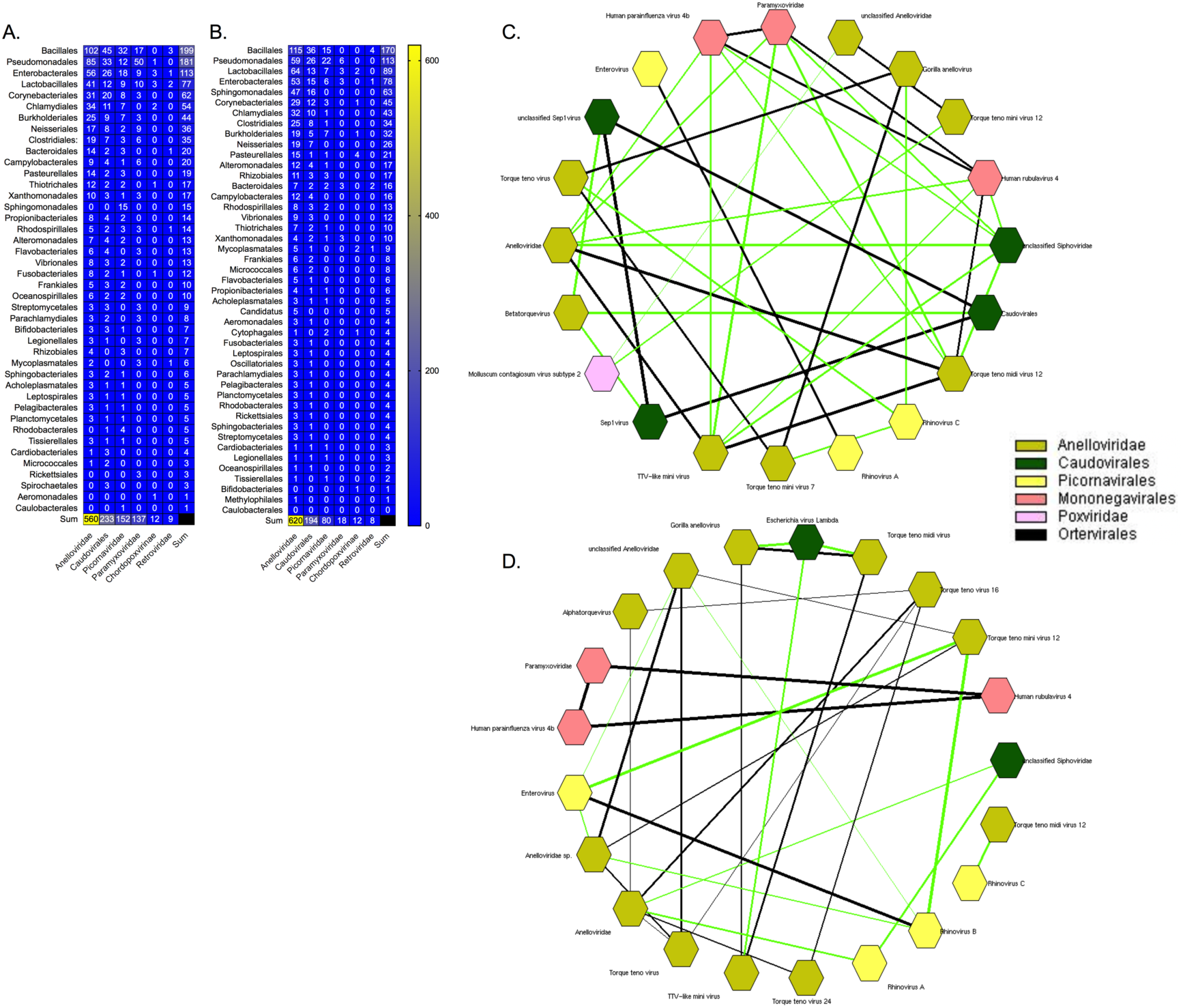
Viral-bacterial and viral-viral co-occurrence pairs. Total number of edges (pairs) between viral families and bacterial orders in the (A) healthy and (B) asthma meta-graphs. Both bacteria and viruses are arranged according to the number of total cross-kingdom connections starting from the one with the highest number (high hubness) towards the one with the lowest (low hubness). Heat map colours; Red: high, Green: low. Virus-virus co-occurrence in the healthy and asthmatic virome: Viral taxa that significantly co-occurred (Pearson p<0.05) are presented as a circular co-occurrence network. All correlations were positive with (C) Pearson R>0.666 in the healthy and (D) R>0.617 in the asthmatic graph. The level of correlation is analogous to the width of the edges of the network. Edges representing correlations amongst taxa from different viral families are highlighted with green. Viral MAGs are colour-coded; Gold: Anelloviridae, Caudovirales: Green, Picornavirales: Light yellow, Mononegavirales: Pink, Poxviridae: Light pink, Black: Ortervirales.

